# Self-similarity and multifractality in human brain activity: a wavelet-based analysis of scale-free brain dynamics^☆^

**DOI:** 10.1101/315853

**Authors:** Daria La Rocca, Nicolas Zilber, Patrice Abry, Virginie van Wassenhove, Philippe Ciuciu

## Abstract

**Background:** The temporal structure of macroscopic brain activity displays both oscillatory and scale-free dynamics. While the functional relevance of neural oscillations has been largely investigated, both the nature and the role of scale-free dynamics in brain processing have been disputed.

**New Method:** Here, we offer a novel method to rigorously enrich the characterization of scale-free brain activity using a robust wavelet-based assessment of self-similarity and multifractality. For this, we analyzed human brain activity recorded with magnetoencephalography (MEG) while participants were at rest or performing a task.

**Results:** First, we report consistent infraslow (from 0.1 to 1.5 Hz) scalefree dynamics (i.e., self-similarity and multifractality) in resting-state and task data. Second, we observed a fronto-occipital gradient of self-similarity reminiscent of the known hierarchy of temporal scales from sensory to higherorder cortices; the anatomical gradient was more pronounced in task than in rest. Third, we observed a significant increase of multifractality during task as compared to rest. Additionally, the decrease in self-similarity and the increase in multifractality from rest to task were negatively correlated in regions involved in the task, suggesting a shift from structured global temporal dynamics in resting-state to locally bursty and non Gaussian scalefree structures during task.

**Comparison with Existing Method(s):** We showed that the wavelet leader based multifractal approach extends power spectrum estimation methods in the way of characterizing finely scale-free brain dynamics.

**Conclusions:** Altogether, our approach provides novel fine-grained characterizations of scale-free dynamics in human brain activity.

**Highlights:**

1. We estimated scale-free human brain dynamics using wavelet-leader formalism.
2. High-to-low self-similarity defined a fronto-occipital gradient.
3. The gradient was enhanced in task compared to resting-state.
4. Scale-free brain dynamics showed multifractal properties.
5. Self-similarity decreased whereas multifractality increased from rest to task.

## 1. Introduction

### 1.1. Scale-free brain activity

Macroscopic brain activity consists of a mixture of synchronized and desynchronized activity [1, 2]. The synchronization of neural oscillations has been hypothesized to mediate neural communication [3, 4, 5, 6], and their coupling, to be involved in information processing [7, 8, 9, 4, 10]. However, the existence, the properties and the functional relevance of scale-free dynamics in brain processing remain an open debate. Scale-free dynamics have been reported in spontaneous brain activity [1] and in data collected with various neuroimaging techniques including fMRI, magnetoencephalography (MEG), electroencephalography (EEG) and local-field-potentials (LFP) [1, 11]. The presence of scale-free dynamics was demonstrated in the infra-slow frequency range (from 0.01 Hz to 1 Hz [1, 12, 13, 14]) and in the slow fluctuations of power of narrow-band neuronal oscillations [15, 16, 17, 18]. The empirical work in both humans and animals has revealed that scale-free dynamics of brain activity were modulated by the levels of wakefulness (*vs.* sleep) [19, 1, 20, 21], consciousness (*vs.* anesthesia) [22, 23], aging and neurodegenerative diseases [24] as well as task performance [25, 1, 26, 27, 17, 18, 28].

The intuition behind the scale-free concept is that the relevant information in the temporal dynamics of a given signal is coded within the relations that tie together temporal scales, rather than in the power of neuronal oscillations in specific bands. Its origin remains however poorly understood. Brain activity recorded with MEG or EEG is most comparable to LFP, and slow dynamic fluctuations likely reflect the up and down states of cortical networks as opposed to spiking activity *per se* [29]. Hence, although fast neuronal activity or avalanches could endogenously produce scale-free infra-slow brain dynamics, a careful statistical assessment remains necessary to draw conclusions on the nature of observed scale-free dynamics [30, 31, 32]. A temporal hierarchy of neural oscillators has been considered a possible source of scale-free brain dynamics [33, 1] as well as the spatial repartition of neural sources. Dendritic filtering [34, 35, 10] or the resistive brain milieu constitute other tentative origins for scale-free dynamics [36]. That the structural configuration of neural networks and their dynamics may be arguably topologically intertwined [37, 35, 38] is also important to keep in mind. To better understand the origins and nature of scale-free brain dynamics, we thus propose to use a rich and robust statistical framework.

### 1.2. Scale-free dynamics modeling and assessment

Scale-free dynamics recorded with neuroimaging techniques have generally been quantified using a 1/*f ^β^* power spectrum model over a large continuum of frequencies. As a result, the empirical assessment has often used Fourier-based spectrum estimation. As an alternative, self-similarity provides a well accepted model for scale-free dynamics that encompasses, formalizes, and enriches the traditional Fourier 1/*f ^β^* spectrum modeling, with models such as fractional Brownian motion (fBm) or fractional Gaussian noise (fGn) [39, 1, 27, 40]. The self-similarity, or Hurst, parameter *H* matches the spectral exponent *β*, as *β* = 2*H* − 1. In the context of brain activity, *H* indexes how well neural activity is temporally structured (via its autocorrelation). Additionally, although *H* has been estimated using Detrended Fluctuation Analysis (DFA) [16, 25, 26, 41, 18, 23], it is now well-documented that wavelet-based estimators provide significant theoretical improvements and practical robustness over DFA, notably by disentangling true scale-free dynamics from non-stationary smooth trends [42, 27, 40].

Often associated with Gaussianity, self-similarity alone does not fully account for scale-free dynamics. The main reason it that self-similarity restricts the description of neural activity to second-order statistics (autocorrelation and Fourier spectrum) and, hence, to additive processes. Yet, it has been proposed that multiplicative processes may provide more appropriate descriptions of neural activity [12]. Independently of, and in addition to selfsimilarity, multifractality provides a framework to model these non-additive processes [43, 24, 44]. Multifractality can be conceived as the signature of multiplicative mechanisms, or as the intricate combination of locally self-similar processes. For instance, if a patch of cortex (i.e. the anatomical resolution of MEG recordings) is composed of several small-networks each characterized by a single self-similar parameter *H*, the multifractality parameter *M* constitutes an index capturing the diversity of *H*s and their interactions within the patch. Qualitatively, the multifractality parameter *M* quantifies the occurrence of transient local burstiness or non Gaussian temporal structures, not accounted for by the autocorrelation function or by the Fourier spectrum (hence, neither by *H* nor *β*). To meaningfully and reliably estimate *M*, it has been theoretically shown that the wavelet-based analysis must be extended to wavelet-leaders [45].

### 1.3. Goals and contributions

The goal of the present work is to produce a rich and reliable characterization of scale-free temporal dynamics in human brain activity, and to provide the field with a robust and reliable procedure to do so. This is made possible (i) by the combined use of self-similarity and multifractality as independent and complementary modeling paradigms, and (ii) by the recourse to the wavelet and wavelet-leader based assessment framework yielding improved performance and robustness to nonstationary trend procedures. The present work investigates the existence and characterization of scale-free dynamics in human cortical activity recorded with MEG, and investigates the modulation of *H* and *M* by resting-state and task.

## 2. Material and Methods

### 2.1. Material

#### 2.1.1. Participants

24 right-handed participants (10 females; mean age of 22.1 ± 1.9 y.o.) took part in the study. All had normal or corrected-to-normal vision and normal hearing and provided a written informed consent prior to the experiment in accordance with the Declaration of Helsinki (2008) and the local Ethics Committee on Human Research at NeuroSpin (Gif-sur-Yvette, France).

#### 2.1.2. Experimental design

The resting-state block lasted 5 minutes during which participants kept their eyes open while staring at a black screen. Participants could mind-wander freely. 5 minutes were selected to be sufficient for an accurate estimation of scale-free properties but not long enough for participants’ cognitive state to drastically change. Resting-state activity was recorded prior to any exposure to task or stimuli. The task block lasted 12 minutes during which participants performed a visual motion coherence discrimination task [46]. In each trial (2.5 s), participants decided which of two intermixed (green and red) clouds of dots was most coherent. Responses were delivered by button press. The experiment was conducted in a darkened soundproof magnetic-shielded room. Participants were seated in upright position under the MEG dewar facing a projection screen placed 90 cm away. The refresh rate of the projector (model PT-D7700E-K, Panasonic Inc, Kadoma, Japan) was 60 Hz. Participants were explained the task and were in contact at all times with the experimenter via a microphone and a video camera. Stimuli were designed using Matlab (R2010a, Mathworks Inc.) with Psychtoolbox-3 [47] on a PC (Windows XP).

#### 2.1.3. MEG data acquisition

Brain activity was recorded in a magnetically shielded room using a 306 MEG system (Neuromag Elekta LTD, Helsinki). MEG recordings were sampled at 2 kHz and band-pass filtered between 0.03 and 600 Hz. Four head position coils (HPI) measured participants’ head position before each block; three fiducial markers (nasion and pre-auricular points) were used for digitization and for alignment with the anatomical MRI (aMRI) acquired immediately after MEG acquisition. Electrooculograms (EOG, horizontal and vertical eye movements) and electrocardiogram (ECG) were simultaneously recorded. Before each experiment, 5 minutes of empty room recordings were acquired for the computation of the noise covariance matrix used in solving the MEG inverse problem.

#### 2.1.4. Anatomical MRI acquisition and segmentation

The *T*_1_ weighted anatomical MRI (aMRI) was recorded using a 3-T Trio MRI scanner (Siemens Erlangen, Germany). Parameters of the sequence were: *FOV* = 256 × 240 × 176 mm^3^, voxel size: 1.0 × 1.0 × 1.1 mm^3^; acquisition time: 7 min46 s; repetition time TR = 2300 ms; inversion time TI= 900 ms; flip angle= 9°; transversal orientation, echo time TE= 2.98 ms and partial Fourier 7/8. Cortical reconstruction and volumetric segmentation of participants’ *T*_1_ weighted aMRI was performed with *Freesurfer*^1^ (RRID: nif-0000-00304). This included: motion correction, average of multiple volumetric *T*_1_ weighted images, removal of non-brain tissue, automated Talairach transformation, intensity normalization, tessellation of the gray-white matter boundary, automated topology correction, and surface deformation following intensity gradients [48]. Once cortical models were complete, deformable procedures could be performed including surface inflation [49] and registration to a spherical atlas [50]. These procedures were adopted using MNE ([51], RRID: scires 000118) to morph individuals’ current source estimates onto the Freesurfer average brain for group analysis.

#### 2.1.5. MEG data preprocessing

Data preprocessing was done in accordance with accepted guidelines for MEG research [52]. Signal Space Separation (SSS) was performed using MaxFilter to remove external magnetic interferences and discard noisy sensors [53]. Ocular and cardiac artifacts (eye blinks and heart beats) were removed using Independent Component Analysis (ICA) on raw signals. ICA was fitted to raw MEG signals, and sources matching the ECG and EOG were automatically found and removed^2^. Then, for the sake of computational efficiency, we downsampled the preprocessed MEG time series at *f_s_* = 400 Hz before applying signal reconstruction following the procedure described in https://github.com/mne-tools/mne-python/blob/master/tutorials/plot_mne_dspm_source_localization.py [51], since scale-free analysis was focused on the low frequency content.

#### 2.1.6. Coregistration and MEG source reconstruction

The co-registration of MEG data with the individual’s aMRI was carried out by realigning the digitized fiducial points with the multimodal markers visible in MRI slices. We used a two-step procedure to ensure reliable MEG-aMRI coregistration: using MRILAB (Neuromag-Elekta LTD, Helsinki), fiducials were aligned manually with the multimodal markers on the MRI slice; an iterative procedure realigned all digitized points (about 30 more supplementary points distributed on the scalp of the subject were digitized) with the scalp of the participant and the MEG coordinates using the mne analyze tools within MNE ([51], RRID:nlx 151346). Individual forward solutions were computed using a 3-layer boundary element model [54] constrained by the individual’s aMRI. Cortical surfaces were extracted with *Freesurfer* (RRID: nif-0000-00304) and decimated to about 5,120 vertices per hemisphere with 4.9 mm spacing. The forward solution, noise and source covariance matrices were used to compute the depth-weighted (parameter = 0.8) minimum-norm estimate [55] inverse operator. The unitless inverse operator was applied using a loose orientation constraint on individuals’ brain data [56] by setting the transverse components of the source covariance matrix to 0.4. Importantly, considering that taking the norm of source dipoles is a nonlinear transformation that may modify scale-free properties [57], we only kept the radial components. Using the individual cortical parcellation provided by *Freesurfer*, reconstructed time series in vertices belonging to the same cortical label (138 labels in total) were grouped and collapsed into a unique time series. In this procedure, the signs of time series within labels were flipped according to anatomical orientation of vertices in such a way that signed activations did not cancel out after averaging (this is a standard label averaging used by the MNE software).

### 2.2. Methods

#### 2.2.1. Scale-free modeling: From Fourier spectrum to selfsimilarity and multifractality

Scale-free dynamics are classically modeled by a power-law decrease of the Fourier power spectrum Γ(*f*) with respect to frequencies *f*: Γ(*f*) ≃*C* |*f*|^−*β*^. Such power laws can be understood as the signatures of the more general and better theoretically framed concept of self-similarity [58]. In essence, self-similarity amounts to modeling scale-free dynamics in data as fractional Gaussian noise (fGn), a Gaussian stationary stochastic process, consisting of the fractional integration (with parameter *H* – 1/2) of a white (i.e., delta-correlated) Gaussian process. The sole parameter *H*, theoretically related to *β* as *β* = 2*H* – 1, governs the entire covariance structure and thus, together with Gaussianity, completely defines temporal dynamics. More precisely, the self-similar parameter *H* quantifies the algebraic decrease of the auto correlation function: *H* = 1/2 indicates the absence of correlation, *H* < 1/2 betrays negative correlation and *H* > 1/2 marks long range positive correlation. While the classical definition of fGn implies 0 < *H* < 1, it can be theoretically extended to *H* > 1 (with the recourse to the notion of generalized processes and tempered distributions [58]), while preserving the original intuition beyond fGn: the larger |*H* − 1/2|, the more structured the temporal dynamics of data (as illustrated in Supplementary Movie 1). Beyond the global control of temporal dynamics via the covariance function, Gaussian self-similarity also implies the absence of fluctuations in the regularity of local temporal dynamics. Such local regularity is often quantified via the Hölder exponent *h*(*t*) *>* 0 [45]. For Gaussian self-similar processes, such as fGn, ∀*t*, *h*(*t*) ≡ *H*.

The multifractal paradigm extends self-similarity by preserving a control of the global temporal dynamics via the covariance function, driven by *H*, while enriching it with possible fluctuations along time of the local regularity *h*(*t*) [45]. Multifractal models, such as multifractal random walk (MRW), are thus essentially stationary non Gaussian processes, defined as the fractional integration (of parameter *H* − 1/2) of a white (i.e., delta-correlated) Gaussian process, whose amplitude is modulated by another independent process, whose covariance decreases logarithmically slowly, with an amplitude controlled by the multifractality parameter *M* ⩾ 0 [59]. Self-similarity parameter *H* preserves the intuitive interpretation of global and overall dependence and structure in the temporal dynamics of data, while the additional multifractal parameter *M* allows local and transient departures from Gaussianity, hence burstiness in temporal dynamics, via fluctuations along time of the local regularity (as illustrated in Supplementary Movie 2).

More technically, multifractal temporal dynamics imply that the fluctuations along time of local regularity are erratic, i.e., the function *h*(*t*) is itself a very irregular function. Therefore, temporal dynamics are not well-described by the local function *h*(*t*), but rather by a global function, the so-called multifractal spectrum 0 ⩽ *D*(*h*) < 1. The multifractal spectrum, which consists of the fractal dimension of the set of points on the real line sharing the same regularity *h*(*t*) = *h* (cf. [45] for a technical definition), thus conveys a global information on the geometrical structure of *h*(*t*), hence on temporal dynamics beyond the mere covariance function. These notions are pedagogically, hence qualitatively, illustrated on synthetic data in Fig. 1.

**Figure 1:**
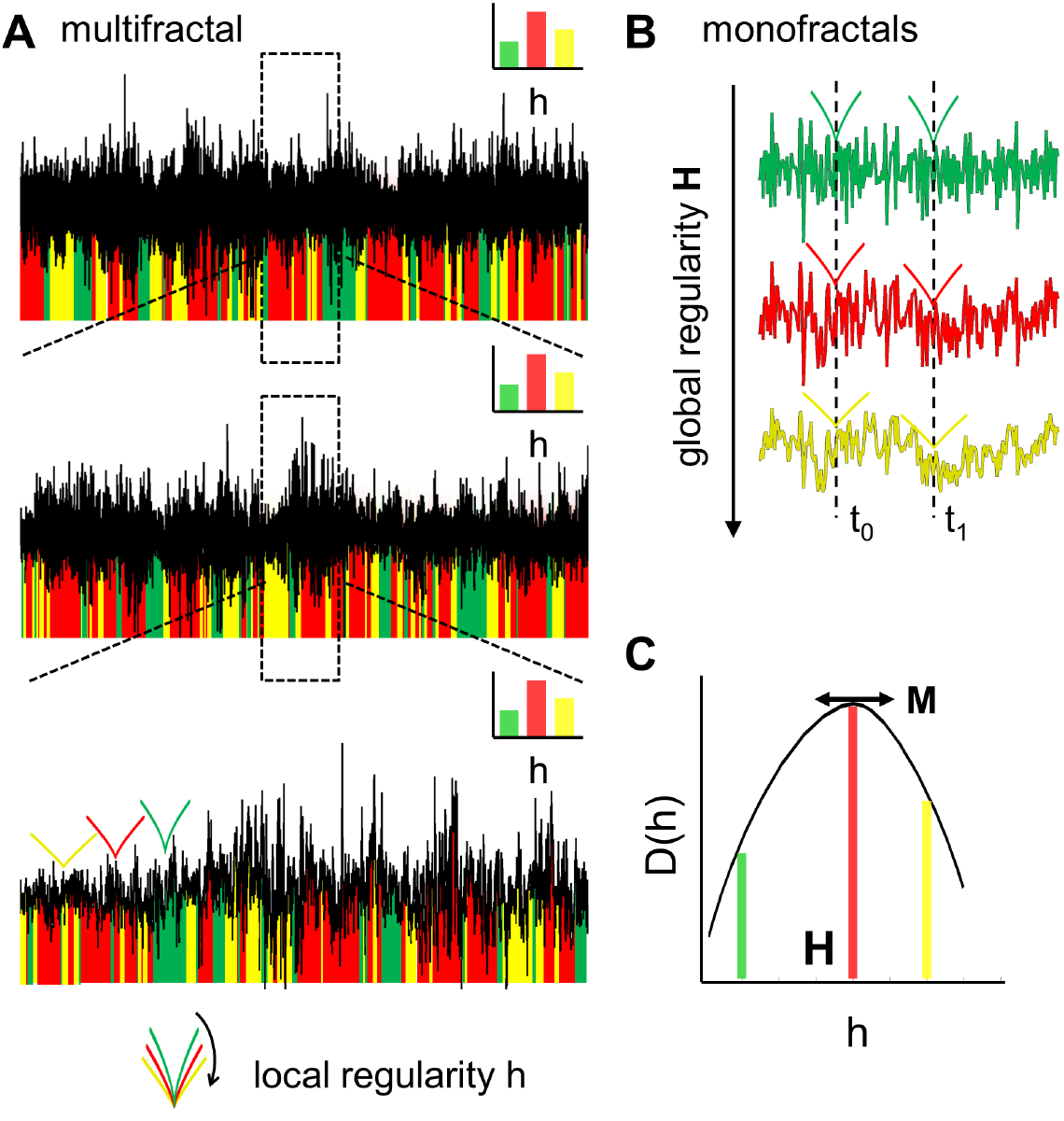
Schematic introduction to Multifractality. (**A**): Multifractal signals observed at three different time scales from coarse (top) to fine (bottom). Local temporal dynamics can be quantified by the Hölder exponent *h*(*t*), a local regularity index. In this pedagogical example, *h*(*t*) can only take three values: red, green and yellow. For multifractal signals, *h*(*t*) is *per se* a very irregular function along time, with all possible *h* existing in any small subpart of the data. (**B**): For monofractal signals (with the same covariance function as the multifractal signals, hence same *H*), no fluctuations of local regularity are observed and the local *h* and the global *H* are everywhere identical. In these three examples of monofractal signals, the global regularity *H* of each signal increases from top to bottom. (**C**): The multifractality illustrated in (**A**) can be captured by a multifractal spectrum *D*(*h*), quantifying by means of fractal (Hausdorff) dimension of the geometrical structure of time points that share the same local regularity *h*(*t*) = *h*. The most frequent Hölder exponent *h*, indicated in red, is closely related to the global self-similarity index *H* whereas multifractality *M* encompasses all local regularities *h* by reflecting the width of *D*(*h*). Importantly, *D*(*h*) remains the same for the whole signal and any subpart, it is hence scale invariant.

While the multifractal spectrum *D*(*h*) can theoretically consist of any shape, it is often efficiently approximated, for practical use, as a parabola controlled by *H* and *M*: *D*(*h*) ≃ 1 − (*h* − *H*)^2^/2*M*. For Gaussian self-similar processes, *M* ≡ 0 and *D*(*h*) = *δ*(*h* − *H*), with *δ* the Dirac-delta function. Parameters *H* and *M* thus provide independent and complementary characterization of scale-free dynamics in data [45], with *M* adding the possibility to model burstiness in temporal dynamics by local departures from Gaussianity, while the global structure of temporal dynamics remains controlled by *H*.

#### 2.2.2. Scale-free analysis: From spectral estimation to wavelet and wavelet leader analysis

The scaling exponent *β* has classically been evaluated by means of spectrum estimation, i.e., by linear regressions in a log-log plot of estimated power spectrum versus frequency (as sketched in Fig. 2). In the present work, all Fourier spectra are estimated using the Welch periodogram procedure. Alternatively, time domain approaches such as detrended fluctuation analysis(DFA) [16], also based on linear regressions, rely on quantifying the power of fluctuations in data increments computed at different lags (acting as scales). It is however now well-documented that multiscale representations, such as wavelet transforms, are well-suited for the analysis of scale-free dynamics and achieve optimal and robust estimation performance cf. e.g., [60, 42].Let *ψ*_0_(*t*) denote a reference pattern, referred to as the mother wavelet, the discrete wavelet coefficients *d_X_*(*j, k*) are defined on a dyadic grid (scale *a* = 2^*j*^ and time *t* = *k*2^*j*^) as: *d_X_*(*j, k*) = ∫*X*(*t*)2^−*j*^*ψ*_0_(2^−*j*^*t* − *k*)*dt*. Under mild conditions on the choice of *ψ*_0_(*t*), it has been shown that for self-similar processes [60]:

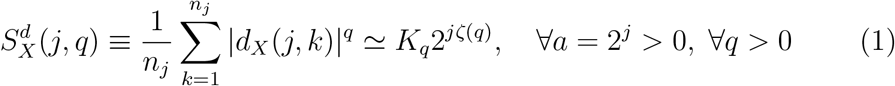

with *ζ*(*q*) = *qH*, thus permitting a robust and efficient estimation of *H* by linear regressions [60]. It can further be shown that, with the particular coice 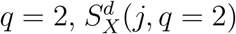, referred to as the wavelet spectrum, can be read as an estimator of the Fourier spectrum Γ(*f*) [42]. Therefore, under elementary transformations, the Fourier and wavelet spectra can be mapped one onto the other, as 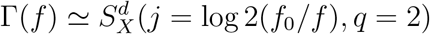, with *f*_0_ a constant that depends on the choice of *ψ*_0_ (and that can be well approximated for a large class of wavelets as *f*_0_ ≃ 3/4 × *f_s_*, with *f_s_* the sampling frequency) [60, 42]. This is quantitatively illustrated in Fig. 2. While both spectra yield equivalent information on the global temporal dynamics, it has been documented that the wavelet spectrum yields a more robust and more reliable estimate of *H* than Fourier spectrum does for *β* [42]. Notably, it was shown that the wavelet spectrum is less prone to bias induced by smooth trends or smooth non stationarity effects, than the classical Fourier spectrum, hence yielding robust estimates of the scale-free exponents.

**Figure 2:**
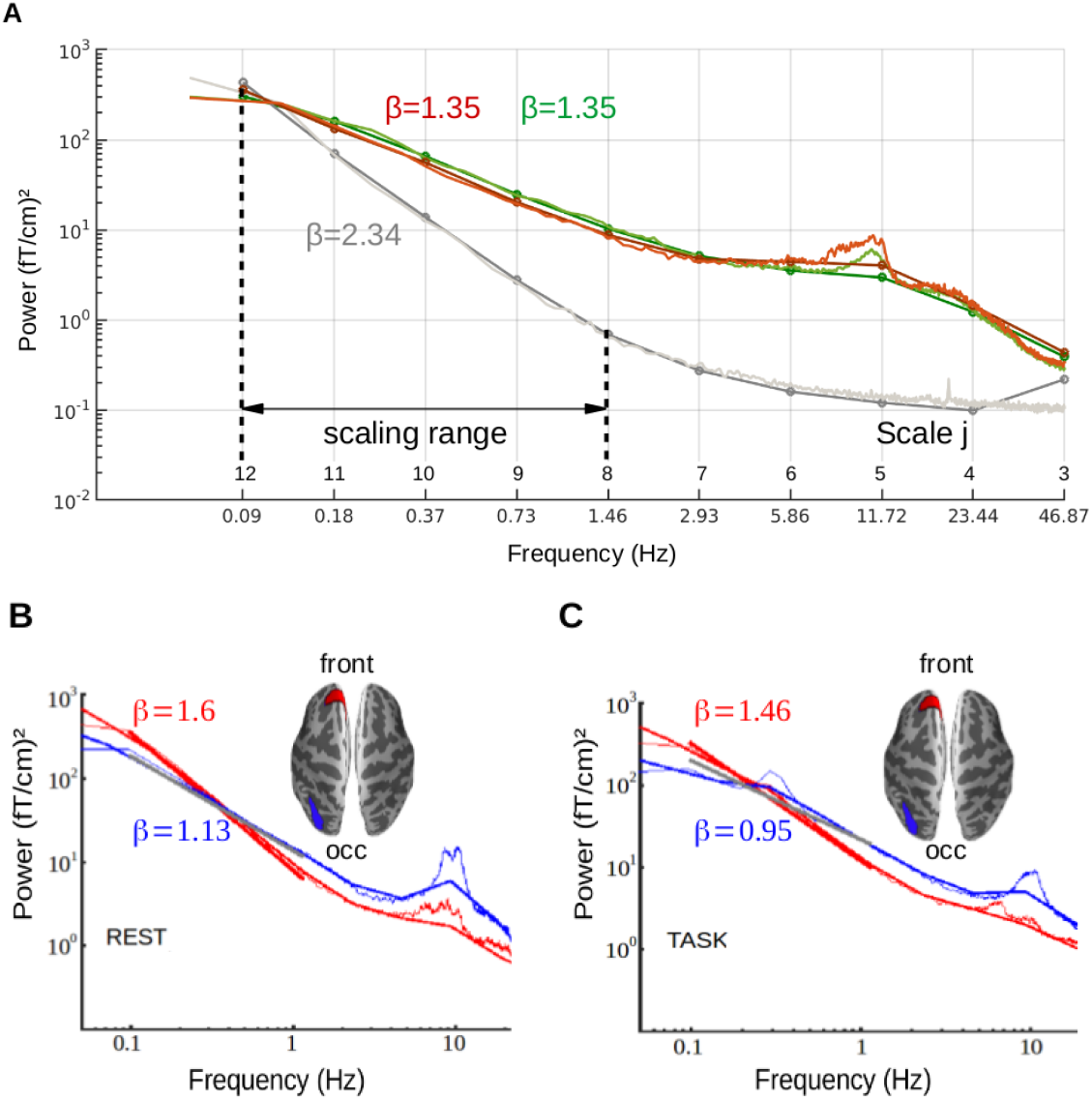
Scale-free brain dynamics: Fourier *vs.* wavelet-based power spectra at rest and during task. (**A**): Group-average Fourier (thin lines) and wavelet (thick lines) power spectra computed in an occipital sensor (inset) for empty room (grey), resting-state (brown) and task (green) recordings. In both Fourier and wavelet log-power spectra, the linear fit indicates scale-free dynamics, and delineates the implicated range of scales (*j* ∈ (8, 12)) corresponding roughly from 0.1 Hz (i.e. *j* = 12) to 1.5 Hz (i.e. *j* = 8). The slopes quantify the scaling exponents *β* (of power spectra 1/*f ^β^*) and the self-similarity index *H*. Human brain activity is characterized by a pink noise (*β* ≃ 1) regime whereas empty room recordings correspond to brown noise (*β* ≃ 2): hence, and importantly, this graph clearly shows that instrumental noise is a not a spurious cause for observing scale-free dynamics in brain activity. (**B-C**): Group-average Fourier (thin lines) and wavelet-based (thick lines) power spectra computed in two frontal (red) and occipital (blue) cortical labels at rest (**B**) and during task (**C**). Larger *β* values correspond to steeper slopes, as shown in the frontal region (**front**, red label) compared to the occipital (**occ**, blue label). All plots clearly show that wavelet and Fourier spectra can be formally mapped one onto the other.

For multifractal processes, when *M* > 0, the scaling exponents *ζ*(*q*) no longer follow the linear form *qH*, but rather consist of a concave function, which in first approximation and for practical purposes can be written as *ζ*(*q*) = *qH Mq*^2^/2. The scaling exponents *ζ*(*q*) are further related to the multifractal spectrum *D*(*h*) via a Legendre transform [45].

For more than one decade [61] it has been proved that a relevant estimate of parameter *M* requires to replace the wavelet coefficients with wavelet-leaders, defined as local suprema of the wavelet coefficients *d_X_*(*j′, k′*), across a local neighborhood *λ_j,k_* = [(*k* − 2)2^*j*^+1, (*k* +1)2^*j*^], for all finer scales 2 *^j′^* ⩽ 2 *^j′^* [45]:

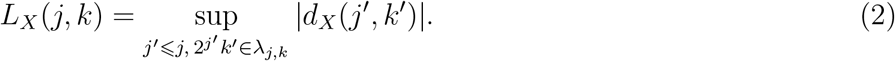

Under mild restrictions, it has been shown that [45]:

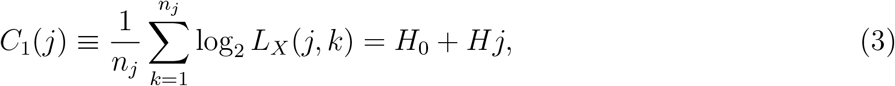

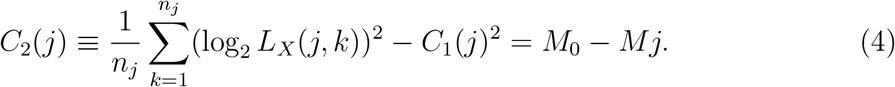

These two fundamental relations show that parameters *H* and *M* can be estimated as linear regressions in diagrams *C*_1_(*j*) vs. *j* and *C*_2_(*j*) vs. *j*, respectively. This is illustrated in Fig. 3. To ease exposition, the functions *C*_1_(*j*) and *C*_2_(*j*) will hereafter be referred to as the wavelet-leader spectra.

**Figure 3:**
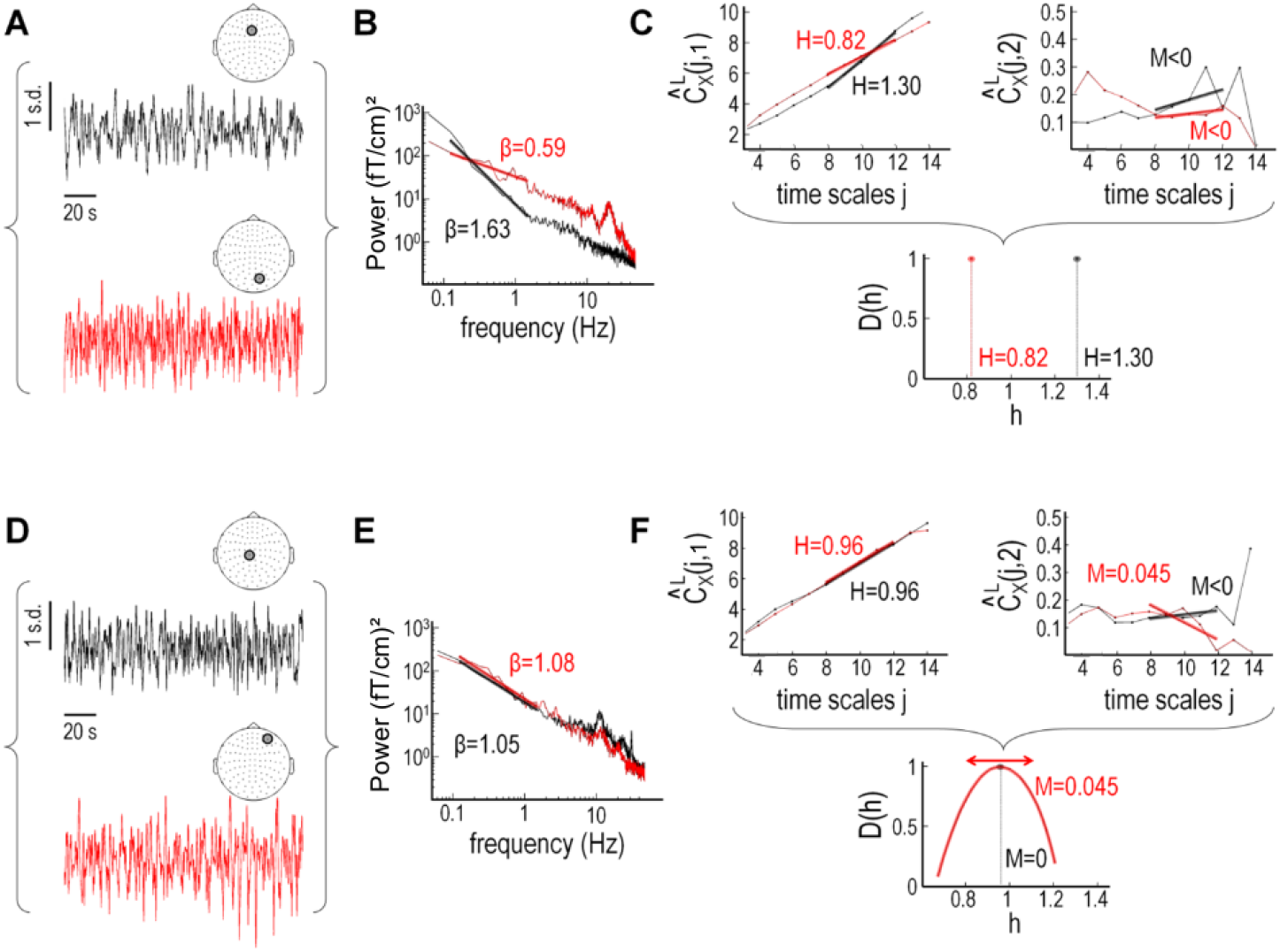
Self-similarity and multifractality in spontaneous infraslow brain activity at rest. Normalized time series representing an individual’s brain activity recorded in two different pairs of MEG sensors at rest (**A-C**) and during task (**D-F**). In (**A-C**), the two signals show a difference in *self-similarity*. In (**D-F**), the two signals show differences in *multifractality* but no differences in self-similarity. The regression analysis was performed over the scaling range (*j*_1_*, j*_2_) = (8, 12) which matched the (0.1, 1.5) Hz frequency range (as a reminder, *j* = 8 corresponds to 1.5 Hz and *j* = 12 corresponds to 0.1 Hz).

## 3. Results: self-similarity and multifractality in human brain activity recorded with MEG

### 3.1. A case study for assessing self-similarity in MEG data: range of frequencies, Fourier vs. wavelet power spectra

Fig. 2 reports the group-average Fourier and wavelet spectra in sensors and in cortical source estimations of the entire MEG data time series. As theoretically expected (cf. Section 2.2.2), the Fourier (thin lines) and the wavelet spectra (thick lines) superimposed very well, yielding consistent patterns across methods. Fig. 2 also shows that both spectra displayed power law behaviors over a broad range of frequencies ranging from roughly 0.1 Hz to 1.5 Hz. Importantly, Fig. 2A compares Fourier and wavelet spectra of human brain MEG data to those of empty-room MEG recordings. This formal comparison unambiguously showed that the spectra differed both in amplitude and in shape: the spectral exponent of human brain recordings was in the so-called pink noise regime (1 ⩽ *β* ⩽ 2) while empty-room recordings rather displayed brown noise temporal dynamics (*β* ⩾ 2). Thus, scale-free dynamics observed in MEG recordings through power spectrum analysis (both Fourier and wavelet) was not caused by instrumental or sensor noise, but rather resulted from macroscopic human brain activity.

Thus, and overall, Figs. 2A-C thus revealed that power law behaviors could be consistently observed during resting-stae and during task, in the range of octaves (*j*_1_, *j*_2_) = (8, 12). This range is associated with frequencies (0.1, 1.5) Hz or, equivalently, with time scales 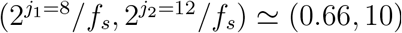 s.

To further provide an intuitive understanding of scale-free dynamics, we compared the Fourier and wavelet spectra of normalized MEG time courses (Fig. 3) recorded from two different sensors in one participant at rest (Fig.3A-C) and during task (Fig. 3.-F). In Fig. 3A, the occipital sensor time series (red) appeared visually less structured over time than the frontal sensor time series (black). This visual appreciation was quantified using a classic Fourier analysis showing a power law behavior at low frequencies (*P* (*f*) ≃ 1/*f ^β^*), with a scaling exponent *β* that was smaller for the occipital sensor (red trace) (Fig. 3B). Conversely, we found that the frontal sensor (black) showed a larger scaling exponent, hence a steeper slope, or equivalently, a stronger temporal autocorrelation, and was quantified by a stronger long-range dependency (Fig. 3B). The same analysis held when conducted from the wavelet spectrum (Fig. 3C), with very satisfactory matches for the estimated scaling exponents *H* and *β* according to *β* = 2*H* − 1.

### 3.2. A case study to go beyond self-similarity in MEG data by assessing multifractality

To demonstrate the interest of a multifractal description in brain activity, we proceeded with multifractal analysis using case study time series. First, using the same example of time series recorded during resting-state as before (Fig. 3A), multifractal analysis yielded negative estimates of the multifractality parameter (Fig. 3C): *M* ⩽ 0 indicating no multifractality at rest. This can be vizualized by the *δ*-shape of the multifractal spectra *D*(*h*), which only differed by their location on the *h*-axis, reflecting different self-similarity exponents *H*. In other words, for these time series, there was no additional information provided by using multifractal analysis.

We then repeated the same analysis on two MEG time-series collected during task (Fig. 3D). These case study time-series where chosen on purpose because they showed very similar Fourier spectra (Fig. 3E), hence displaying the same *β*s, and predictably, the same held true for the wavelet spectra (Fig. 3F) showing the same *H*s. Interestingly however, the frontal signal (red) appeared far more irregular and locally bursty than the central one (black). This was again quantified using multifractal analysis (Fig. 3F), which revealed that although *H* (top left) was identical in both frontal and central time series, the frontal time series was characterized by a positive *M* = 0.045 > 0 (hence, displayed multifractality), while the central sensor did not (*M* < 0). The multifractal spectra for both time series thus summarized the two case study observations: whereas the location of their peaks coincided (same *H*), only one spectrum (red) showed a large parabola shape (*M* > 0).

These examples were chosen as pedagogical illustrations of the potential richness of scale-free temporal dynamics found in brain time series. Specifically, while the typical power spectrum analysis would conclude that these different time series share the same scale-free characteristics, multifractal analysis clearly showed differences in their temporal dynamics by quantifying the existence of transient and local irregularities observed in the frontal sensor (red) that did not exist in the central sensor data (black). Multifractality thus complements self-similarity in the characterization of scale-free dynamics in time series by quantifying local transient dynamics that are not well accounted for by the autocorrelation or by the Fourier spectrum.

### 3.3. Group-level analysis of scale-free brain dynamics

Having extended the framework for the assessment of scale-free brain dynamics to self-similarity *H* and multifractality *M*, we then proceeded with a comprehensive analysis of scale-free brain activity across all individuals (*n* = 24). For this, we assessed scale free activity in source reconstructed time series averaged within each cortical regions (see averaged within each cortical Section 2.1.6). The estimation of parameters *H* and *M* relied on the wavelet-leader multifractal formalism described in Section 2.2.2: *i.e.*, MEG wavelet-leader spectra *C*_1_(*j*) and *C*_2_(*j*) were systematically computed on resting-state and task recordings separately for each cortical label and on *a per* individual basis. Results were then averaged across individuals to form *C̅*_1_(*j*) and *C̅*_2_(*j*), respectively. Importantly, since linear regression and group-level averaging were both linear after taking log in Eqs. (3)-(4), we could interchange them without impacting the results. For this reason, in what follows, we illustrated group-level values of *C̅*_1_(*j*) and *C̅*_2_(*j*) in log-scale diagrams from which we deduced the respective group-level *H* and *M*. The latter actually matched the group-level averages of subject-specific values of *H* and *M*, shown in the cortical maps of Figs. 4-5.

### 3.4. A fronto-occipital gradient of self-similarity

Fig. 4A reports the grand average *C̅*_1_(*j*) for resting-state obtained in two cortical labels (one frontal in red, one occipital in blue). The self-similarity exponent *H* was found to be larger in the frontal label as compared to the occipital label. To systematically quantify this effect, the calculation of *C̅*_1_(*j*) was conducted over the whole cortical surface. Using T-statistics, the null hypothesis *H* = 0.5 was tested at the group level. To account for multiple comparisons across the 138 labels covering the whole cortical surface, a correction was implemented using the false discovery rate (FDR) detection at *α* = 0.05: *p*_corrected_ < 0.05. Fig. 4C reports the spatial distribution of statistically significant mean values of *H* (*H* > 0.5), yielding a key finding: the spatial distribution of estimated *H*s during rest revealed a fronto-occipital gradient, in which *H* significantly decreased from frontal (*H* ≃ 1.2) to occipital regions (*H* ≃ 0.8 − 0.9). This gradient was consistent with prior observations of scale-free activity observed in MEG and EEG recordings [36]: larger *H* in frontal regions (i.e., steeper slopes for the spectra) would indicate stronger and longer temporal correlations,*i.e.* more-structured temporal dynamics, compared to occipital regions.

**Figure 4:**
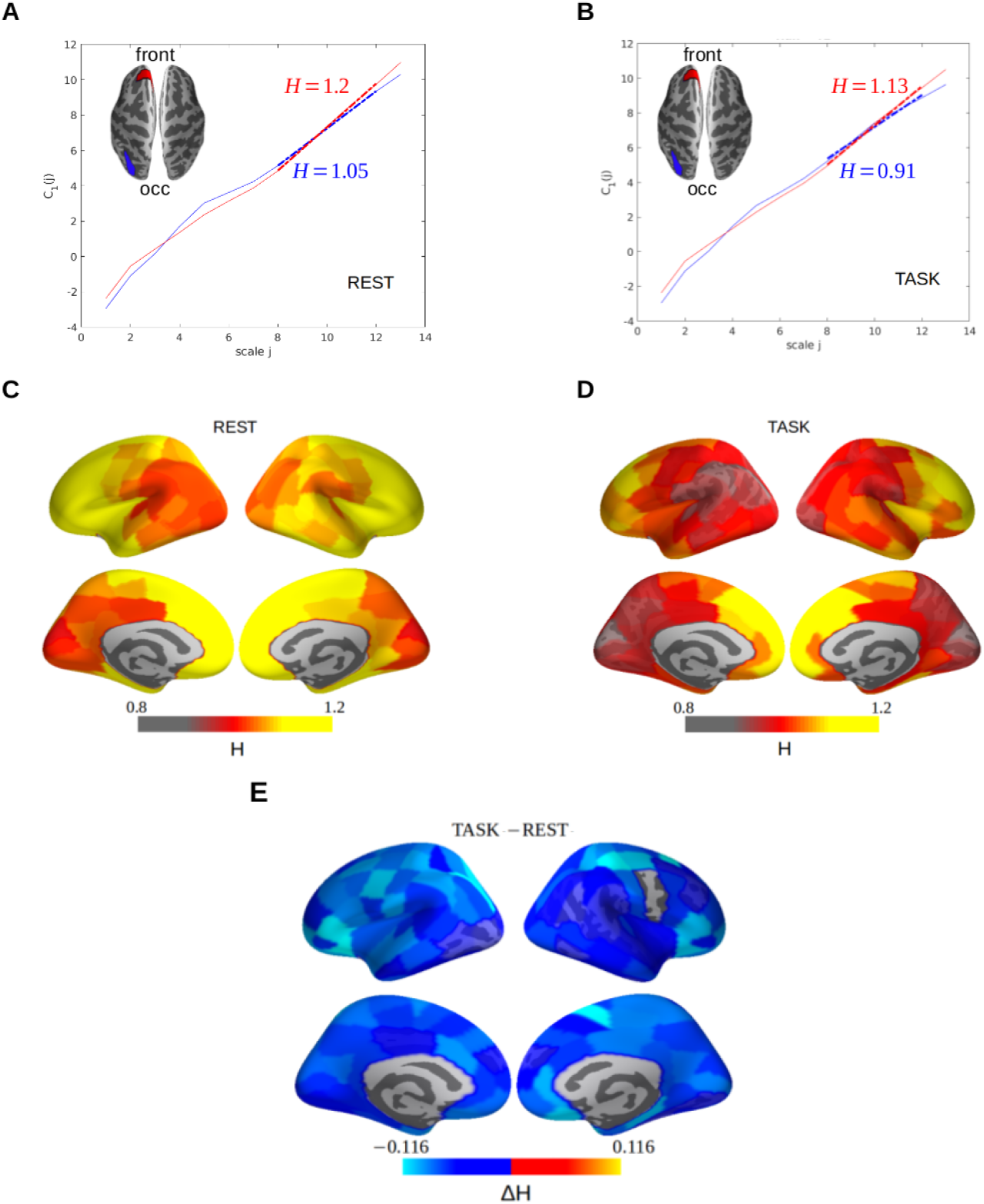
Fronto-occipital gradient of self-similarity. For comparison with Fig. 2, we show group-average wavelet-leader structure functions *C̅*_1_(*j*) in the same frontal (red) and occipital (blue) cortical labels during rest (**A**) and task (**B**) blocks. The linear fits were computed over the scaling range 8 ⩽ *j* ⩽ 12 and matched the (0.1, 1.5) Hz frequency range used before for linear regression in the power spectra. The associated slopes provides estimates of group-level Hurst exponents *H*. (**C**-**D**): Group-average cortical maps (lateral and medial views on top and bottom, respectively, left hemisphere on the left) of Hurst exponents *H* at rest and during task, respectively. In both rest and task, a fronto-occipital gradient of self-similarity could be observed going from higher *H* in frontal regions to lower *H* in parieto-occipital regions). (**E**): Cortical maps contrasting *H* in task and resting-state testing the null hypothess that *H*_TASK_ = *H*_REST_. The statistical significance was assessed on a per label basis by computing a paired Student t-test and correcting for multiple comparisons with FDR at *α* = 0.05. Estimates of *H* were smaller in task than in rest as shown by negative differences (∆*H* = *H*_TASK_ – *H*_REST_ *<* 0). This contrast indicated that, globally, self-similarity significantly decreased when participants performed a task as compared to when they rested.

### 3.5. During task, an overall decrease of self-similarity accentuates the fronto-occipital gradient

In Fig. 4D, the spatial distribution of *H* significantly departed away from 0.5 during task yielding, by comparison to rest, another key finding: the decrease of *H* during task appeared to be global and almost significant everywhere over the cortical surface. Interestingly, the anatomical fronto-occipital gradient at rest appeared to be further strengthened during perceptual task completion (cf. lateral views in Fig. 4D). We contrasted the *H* parameter estimates between rest and task using paired *t*-tests. FDR was applied to correct for multiple comparisons across cortical labels at *α* = 0.05. In Fig. 4E, the statistical assessment of changes in *H* between rest and task confirmed our qualitative appreciation. Specifically, *H* was significantly diminished during task in numerous cortical regions including occipital, parietal, and primary motor cortices as well as right supplementary area (SMA) and ventrolateral prefrontal cortex (vlPFC) bilaterally. All these regions were previously shown to be essential in the perceptual task participants were engaged in [46]. Conversely, cortical regions belonging to the default mode network or DMN (*e.g.*, the right medial prefrontal cortex) showed a smaller decrease of *H* by less than 10 % (decrease from 1.2 to 1.12) during task. This may indicate that DMN regions implicated in monitoring resting-state may show reduced to opposite trends than regions implicated in task performance.

### 3.6. Weak multifractality in resting-state and default mode network (DMN)

The group-average *C̅*_2_(*j*) at rest for two cortical labels is illustrated in Fig. 5A. We found no multifractality (*M* < 0) in the frontal label (red) but found multifractality (*M* = 0.017) in the occipital label (blue). As previously done for estimates of *H*, we performed the analysis of multifractality at rest over the whole cortical surface. Using T-statistics, the null hypothesis *M* = 0 was tested at the group-level. The same FDR correction at *α* = 0.05 was applied to correct for multiple comparisons across labels. Fig. 5C reports the spatial distribution of statistically significant mean values of *M* (*M* > 0). At rest, the presence of multifractality was confined to a few regions: the posterior superior temporal sulci, the occipital cortex, the right temporo-parietal junction and the frontal cortices, bilaterally. Additionally, multifractality was observed in two additional regions of the Default Mode Network (DMN) lateralized to the right hemisphere, namely the posterior cingulate cortex and the middle prefrontal cortex. The observed values of *M* mostly ranged between 0.01 and 0.02, with the exception of the frontal poles which reached *M* = 0.03.

**Figure 5:**
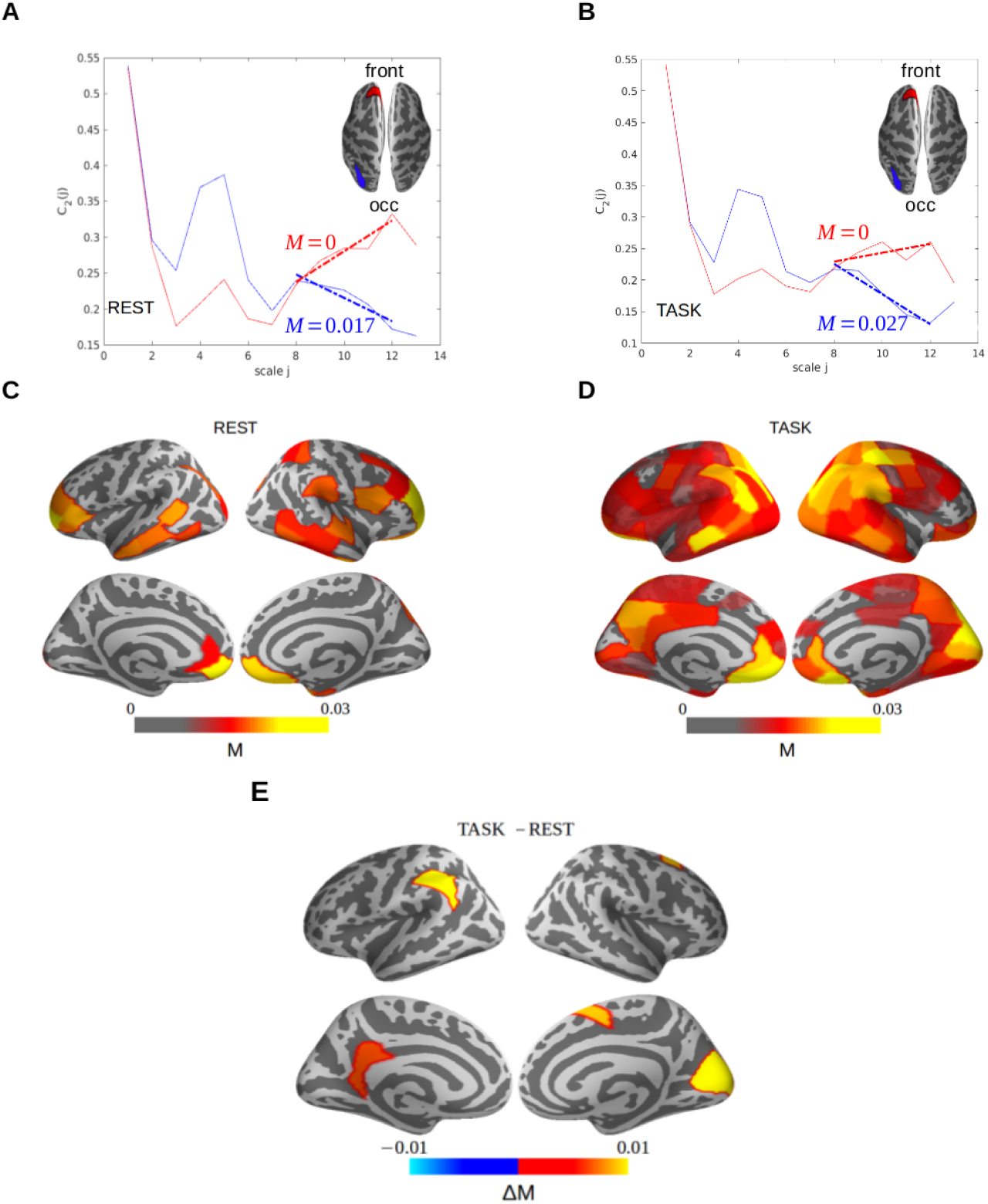
Multifractal brain activity. Group-average wavelet-leader structure functions *C̅*_2_(*j*) in the same frontal (red) and occipital (blue) labels at rest (**A**) and during task (**B**). The linear fits were computed over the scaling range 8 ⩽ *j* ⩽ 12 and matched the (0.1, 1.5) Hz frequency range used before. The associated slopes provided estimates of the multifractal exponents *M*. (**C**-**D**): Statistically significant (*H*_0_: *M* = 0) grand-average cortical maps of multifractal exponents *M* at rest and during task, showing sparser topographies than for self-similarity *H*, especially at rest. (**E**): Cortical maps contrasting *M* in task and resting-state testing the null hypothesis that *M*_TASK_ = *M*_REST_. The statistical significance was assessed on a per label basis by computing a paired Student t-test and correcting for multiple comparisons with FDR at *α* = 0.05. Estimates of *M* were bigger in task than in rest as shown by positive differences (∆*M* = *M*_TASK_ *M*_REST_ *<* 0) in several regions involved in the task, notably visual, parietal and motor cortices. This contrast indicated that, locally, multifractality significantly increased when participants performed a task as compared to when they rested.

### 3.7. Multifractality: localized increase in regions engaged in the task

During task, the group-level wavelet-leader spectra *C̅*_2_(*j*) (Fig. 5B) suggested an absence of multifractality in frontal regions (*M* = 0) but an increase of multifractality in occipital regions. *M* was increased by about 60 % in occipital cortices, hence showing steeper slopes for *C̅*_2_(*j*). A statistical assessment of multifractality over the whole cortical surface during task revealed a spatially extended set of cortical regions showing significant multifractality (Fig. 5D; mean values of *M* (*M* > 0)). Additionally, the multifractal parameter *M* took overall larger values compared to the distribution we had observed during resting-state. We notably found larger *M* values in cortical regions involved in the perceptual task participants were engaged in [46], namely: visual cortices (primary, secondary and visual motion region (hMT+)) as well as parietal cortices and the posterior superior temporal sulci.

When statistically contrasting the *M* parameter estimates between rest and task (paired *t*-tests, FDR correction for multiple comparisons across cortical labels at *α* = 0.05), we found significant changes in *M* in brain regions including the right occipital cortices, SMA, the left temporo-parietal junction and posterior cingulate cortex (Fig. 5E). These changes corresponded to an increase in multifractality, though they remained limited in magnitude to a maximal increase of 0.01. This was a remarkable observation considering that the analysis was conducted over the whole cortical surface with no *a priori* restriction on the timing of the stimuli or cognitive operations implicated in the decision-making; rather, our analysis was performed over the whole time series.

Altogether, these results suggest that multifractal characterization of brain activity may capture relevant signatures of brain processing that are associated with task-relevant brain regions.

### 3.8. Covariation of self-similarity and multifractality from rest to task

So far, we reported significant differences for both *H* and *M* when contrasting rest and task, namely: while self-similarity *H* significantly decreased in task as compared to rest (Fig. 4E), multifractality *M* significantly increased in task as compared to rest (Fig. 5E). Additionally, the changes in *M* were confined to a limited number of brain regions whereas the changes observed in *H* were more global, thereby yielding a global accentuation of the fronto-occipital gradient. Considering the possible overlap of cortical regions displaying both changes, we then asked to which extent the two characteristics of scale-free dynamics may be related. We correlated the changes in *H* and *M* from rest to task on a label-by-label basis (Fig. 6). This analysis revealed that in some of the cortical regions showing task-related multifractality (Fig. 6B), there was a significant negative correlation between individual changes from rest to task of *H* (∆*H* = *H*_TASK_ − *H*_REST_) and *M* (∆*M* = *M*_TASK_ − *M*_REST_) (Fig. 6B).

**Figure 6:**
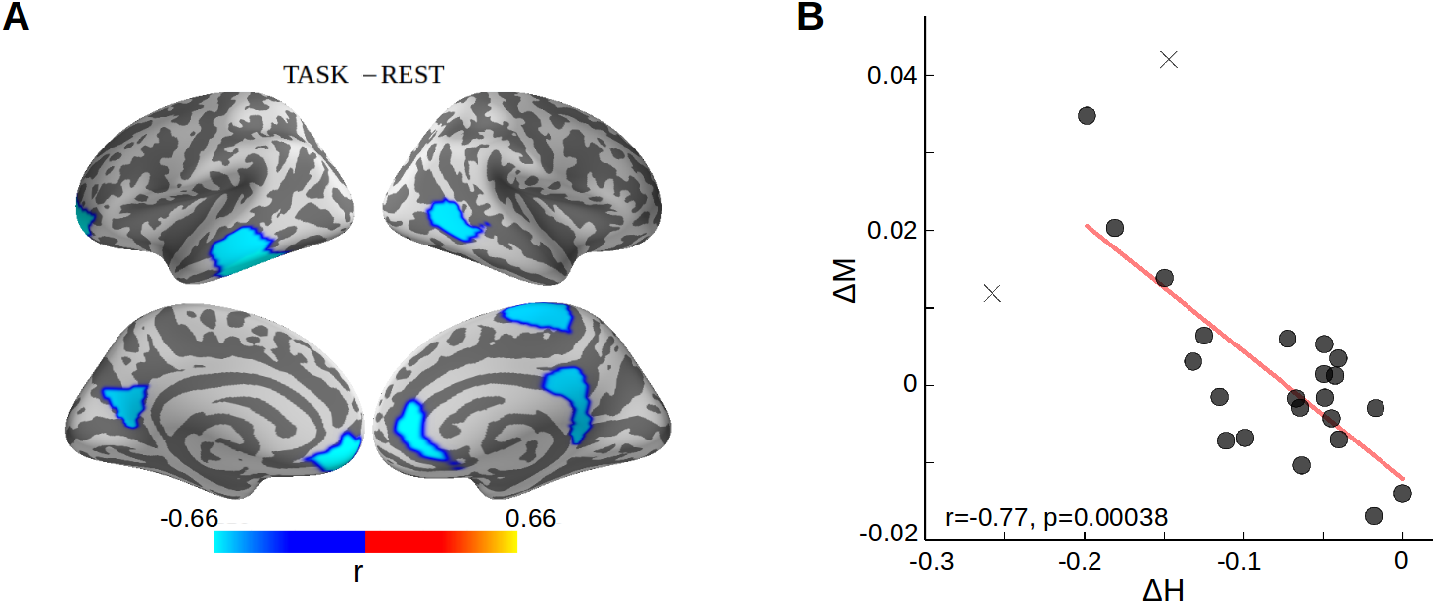
Covariations of self-similarity and multifractality. Significant negative correlation between changes of *H* (∆*H* = *H*_TASK_ − *H*_REST_) and *M* (∆*M* = *M*_TASK_ − *M*_REST_). (**A**): Cortical source estimates associated with significant correlations (color-coded Pearson’s r values) observed between ∆*H* and ∆*M*. The null hypothesis reads *r* = 0 in each cortical label. Statistical significance was assessed on a per individual and per label basis by computing a paired Student t-test. Corrections for multiple comparisons were performed using FDR at *α* = 0.05. (**B**): Scatter plot of ∆*H* versus ∆*M*, averaged over all brain regions reported in (**A**). The significant negative correlation indicated concomitant local decreases of *H* (negative ∆*H*) and increase of *M* (positive ∆*M*). Each dots is an individual and crosses are outliers (*n* = 24).

These results constituted a particularly important finding: theoretically, *H* and *M* are independent parameters, which model very different aspects of scale-free dynamics. While self-similarity *H* provides insights on the temporal autocorrelation of brain activity, *M* informs on the burstiness of the signals. The observed negative covariation between self-similarity and multifractality is non-trivial and crucially suggests a potential coupling in the covariation of both indices. We discussed these findings further below.

## 4. Discussion

To briefly sum up, our key findings are:the existence of a fronto-occipital gradient of self-similarity in the human brain which increases during task as compared to rest. Second, and to the best of our knowledge, we observed for the first time multifractality on MEG data collected in a healthy human population and describe an anatomical distribution of multifractality during resting-state and task. Local changes in multifractality in task as compared to rest indicate a possible functional relevance of multifractal infra-slow dynamics in brain processing. Our empirical results raise several points of discussions and conclusions regarding the assessment of scale-free temporal brain dynamics. We discuss the main ones below.

### 4.1. Robust description of infraslow, scale-free dynamics, in human brain activity

Overall, our results support the notion that scale-free temporal dynamics constitute a signature of human brain activity as recorded with MEG. We showed that scale-free properties were neither induced by, nor to be confused with, instrumental or sensor noise considering that empty-room recordings did not display the same characteristics. Scale-free dynamics were observed in a range of frequencies corresponding to 0.1 ⩽ *f* ⩽ 1.5 Hz. Such time scales are consistent with currently available data in the literature for the estimations of *H* or *β* [17, 1, 13, 14], and typically characterize infra-slow neural dynamics [6, 12, 13]. Scale-free temporal dynamics conceptually implies that, within the scaling range, no frequency plays a particular role. Conversely, all frequencies in that range contribute jointly and in a related manner to the described dynamics. Such relation is quantified by *β*, or in a richer framework which we propose here, by the joint description of *H* and *M* parameters. Scale-free temporal dynamics thus correspond to arrhythmic signatures confined to infra-slow brain dynamics, which complement oscillatory activity typically seen at higher frequencies (> 2 Hz) [6, 12, 13]. It is also noteworthy that while the concept of scale-free temporal dynamics theoretically implies the absence of any specific time scale, in practice, scale-free analyses have covered a finite range of frequencies, 0.1 ⩽ *f* ⩽ 1.5 Hz.

We thus propose that the classical parameter *β* used to model scale-free dynamics as a power-law decay of the Fourier spectrum can be efficiently replaced by the self-similarity parameter *H*, which models the decay of the wavelet spectrum. While both exponents are theoretically equivalent and related (*β* = 2*H* − 1), *H* and wavelet analysis benefit from improved estimation performance (robustness to smooth non-stationarities). The larger *H* (or *β*) - *i.e.*, the steeper the decay of power laws -, the more structured the temporal dynamics of the time series -*i.e.,* the stronger the long range dependency quantified by the temporal correlations.

### 4.2. Multifractality: going beyond self-similarity

In addition to self-similarity, our results demonstrated the existence of multifractality while participants performed a task and, to a lesser extent, during resting-state. This specific result underline the richness of infraslow brain dynamics and of the usefulness of the framework we propose to characterize scale-free brain activity. Specifically, we showed that multifractality allows distinguishing time series that share the same global correlation structure (*i.e.*, the same self-similarity) but different local transient structures and burstiness over time (*i.e.*, multifractality). In other words, multifractality quantifies local scale-free temporal dynamics as transient departures from Gaussianity and support the recent mention that multiplicative processes should be taken into account in the assesment of macroscopic brain activity [12]. Hence, our approach is key to proper scale-free modeling although it remains seldom discussed in the neuroscience literature [43, 24, 27]. Additionally, and from a signal processing perspective, our results suggest that the multifractal random walk is a likely more accurate modeling than the fractional Brownian motion to describe spontaneous brain activity in the infraslow regime (< 2 Hz).

### 4.3. Anatomical distribution of self-similarity and multifractality in resting-state activity

The spatial distributions of self-similarity and multifractality quantified at rest and during task were obtained using the theoretically robust and practically efficient wavelet-leader multifractal framework [45]. With this approach, we observed a fronto-occipital gradient of the self-similarity parameter *H* in resting-state. This observation was congruent with previous findings in the literature [36, 14], but also extended them from scalp level to cortical source estimates. The fronto-occipital gradient corresponded to larger values of self-similarity in frontal regions and lower values in posterior regions. This pattern converges with the known distribution of temporal scales at which neural processing operate: a recent meta-analysis has notably showed a hierarchy of intrinsic time-scales going from slower dynamics in frontal to faster dynamics in sensory cortices [62]. Comparable temporal hierarchies have been functionally described in the human visual system [63] and across brain systems [64]. These temporal hierarchies are functionally compatible with finer time scales needed for sensory sampling, and integrative processes over longer time scales occurring in frontal cortices for higher cognitive operations [65, 66, 67]. By indexing the anatomical distribution of temporal autocorrelation functions, the fronto-occipital gradient in *H* provides an alternative means to characterize the hierarchy of temporal scales in cortex.

Additionally, during resting-state, the presence of weak multifractality localized in regions of the DMN (fronto-polar, middle prefrontal cortex and the PCC) was supplemented by multifractal spontaneous brain activity in the occipital cortex and along the superior temporal sulcus. This weak multifractality, naturally one order smaller than values of *H*, is consistent with well-behaved multifractal synthetic models [45] or values reported for brain data [43, 68, 24, 69, 19, 27, 70]. The presence of *M* in DMN and beyond suggested that multifractality may be a relevant index for brain processing.

### 4.4. Global decrease of self-similarity from rest to task

By contrasting brain activity during engagement in a task against resting-state, we observed a general decrease of *H* over the whole cortex, suggesting an overall and global shortening of temporal autocorrelation during task performance. Additionally, the decrease in self-similarity was not uniform across brain regions, which contributed to the strengthening of the fronto-occipital gradient. In other words, relatively less short-time dynamics were found in frontal regions and more short-time dynamics were observed in posterior regions during task than during rest. The accentuation of the fronto-occipital gradient in *H* between rest and task is overall consistent with faster and richer dynamics deployed for the analysis of sensory information in cortical regions engaged in the task [71]. This observation also converges with previous fMRI studies showing a lower regional *H* during task than during resting-state [26, 27] and stronger decreases of *H* with higher cognitive loads [72]. The most salient differences of self-similarity were observed in regions involved in the task (occipital cortex, motor cortex, SMA and vlPFC), *i.e.* the decrease in *H* was the largest in these regions. This observation is in line with the hypothesis that self-similarity may quantify neural excitability, with smaller values of self-similarity indexing higher levels of neuronal excitability in a given brain region [1, 26, 71, 18].

### 4.5. Local increase of multifractality from rest to task

Although we found a large number of cortical labels showing a significant presence of multifractality during task, contrasting task against rest revealed increases of *M* in only a small number of cortical regions. The relatively small changes of *M* in magnitude, the limited sample size (*i.e.* 24 individuals only) and the potentially large inter-individual variability may explain why only a fraction of cortical regions were reported as statistically significant in the paired *t*-test. Nevertheless, the presence of the highest *M* values in regions (occipito-parietal cortices, visual motion area, pSTS) involved in the visual motion discrimination task used here [46] suggests that multifractality may be functionaly relevant to cortical processing. The local changes of multifractality would be consistent with the notion that multifractality may reflect the combination of multiplexed self-similar processes,*i.e.* the superimposition of several self-similar processes associated with different neural populations within the same cortical patch (given the limits of the spatial resolution with MEG). As such, one working hypothesis for multifractality in brain processes is that it may index the number of neural processes within a cortical region employed in a given task. This working hypothesis will be actively investigated.

### 4.6. Covariation of self-similarity and multifractality from rest to task

We evidence an interesting covariation pattern in self-similarity and multifractality from rest to task: the MEG brain dynamics evolved from well structured and long term correlated global temporal dynamics (large *H*) with weak burstiness (*M* ≃ 0, weak multifractality) at rest, to less structured global temporal dynamics (lower *H*, lesser long range dependence, or more power at the upper bound of the scaling range, *i.e.,* around 1Hz) during task performance, showing though much larger transient irregular and non Gaussian behaviors (larger *M*, multifractality). Let us emphasize that this covariation (decrease in *H*, increase in *M*) was non trivial and was not induced by the modeling nor by the analysis we undertook. This covariation thus constitutes a signature of the changes induced in brain dynamics when participants engaged in a perceptual discrmination task.

Our tentative explanation for this covariation is the following: the local decrease of temporal autocorrelation (*H*) suggests that neural populations in a given cortical region *and* at a large temporal scale (lower infra-slow *i.e.* ≃ 10 s) tend to operate more independently while, at the same time, the increase of temporal burstiness (*M*) in the same region suggests that the same neural populations may interact at finer temporal scales (higher infra-slow, *i.e.* ≃ 1s). Distinct dynamic modes may thus take place as a function of task requirements: while neural excitability may be sufficient to detect the presence/absence of a stimulus in the environment [1, 26, 17], temporal multiplexing may be required for thorough analysis of sensory inputs. In other words, temporal multiplexing may occur when a certain level of neural excitability has been reached.

## 5. Conclusions

Relying on the robust and efficient wavelet and wavelet-leader analysis framework, our present contribution showed that multifractality provides a fruitful paradigm to complement self-similarity in the modeling of scale-free temporal dynamics and infraslow macroscopic brain activity. We showed that spontaneous human brain activity at rest is well characterized by a a strong self-similarity and weak multifractality, indicating a significantly globally-structured activity, with long range dependencies. The strength of this structured activity showed a fronto-occipital gradient. We showed that performing a task induced a non trivial (negatively correlated) local coupling of self-similarity and mutifractality with an overall decrease of self-similarity (yet, strengthening of the fronto-occipital gradient) accompanied by a local increase of multifractality in task-relevant brain regions. Overall, this pattern indicates less structured (or less correlated) temporal dynamics yet bursty occurences of well-structured local scale-free patterns (not accounted for by self-similarity but well quantified by multifractality). Altogether, these observations support the hypothesis that (i) self-similarity, as indexed by parameter *H*, inversely reflects neural excitability, with large *H* corresponding to lower excitability and vice versa and that (ii) multifractality, indexed by *M*, might code for multiplexing of neural processes.

The present analysis of scale-free dynamics in brain temporal dynamics will be continued by exploring the benefits of using more refined analysis tools based on *p*-leaders [73] or on multivariate models, rather than univariate, for self-similarity [74] and multifractality [75, 70].

## Footnotes

1 http://surfer.nmr.mgh.harvard.edu

2 https://github.com/mne-tools/mne-python/blob/master/tutorials/plot_artifacts_correction_ica.py

